# Low dose inocula of SARS-CoV-2 B.1.1.7 variant initiate more robust infections in the upper respiratory tract of hamsters than earlier D614G variants

**DOI:** 10.1101/2021.04.19.440414

**Authors:** Bobo Wing-Yee Mok, Honglian Liu, Siu-Ying Lau, Shaofeng Deng, Siwen Liu, Rachel Chun-Yee Tam, Timothy Ting-Leung Ng, Jake Siu-Lun Leung, Pui Wang, Kelvin Kai-Wang To, Jasper Fuk-Woo Chan, Kwok-Hung Chan, Kwok-Yung Yuen, Gilman Kit-Hang Siu, Honglin Chen

## Abstract

There is a lack of experimental evidence to explain how the B.1.1.7 variant spreads more quickly than pre-existing variants in humans. We found that B.1.1.7 displays increased competitive fitness over earlier D614G lineages in an *in-vitro* system. Furthermore,, we demonstrated that B.1.1.7 variant is able to replicate and shed more efficiently in the nasal cavity than other variants with lower dose and shorter duration of exposure.

---

In late 2020, a novel SARS-CoV-2 variant of concern (VOC), VOC 202012/01 (lineage B.1.1.7) was identified in the United Kingdom. This B.1.1.7 variant containing multiple mutations in spike^1^ has become dominant in the UK and is now rapidly spreading across multiple countries^2^. It is thought that this VOC has the potential to spread more quickly and with higher mortality than the pandemic to date^3^. Recently, using multiple behavioural and epidemiological data sources, Davies et al. estimated that the VOC 202012/01 variant (lineage B.1.1.7) has a 43–90% higher reproduction number than pre-existing variants in England^4^. In another study, Davies et al. indicated that among specimens collected in the UK in early 2021, higher concentrations of virus were found on nasopharyngeal swabs from B.1.1.7 infected individuals, as measured by Ct values from PCR testing^5^. However, there is a lack of experimental evidence to support the expectation that B.1.1.7 does indeed spread more quickly than pre-existing variants.

The B.1.1.7 variant of SARS-CoV-2 harbours 21 non-synonymous point mutations and 3 deletions in comparison to the reference genome (accession number: NC_0.45512.2). Of these, 8 mutations and 2 deletions are involved in changes in the spike protein, which interacts with the host cell receptor, angiotensin-converting enzyme 2 (ACE2), and mediates virus entry into host cells^6^. These spike mutations include the deletion ΔH69/ΔV70, which has arisen in multiple independent lineages and is suggested to associate with increased infectivity and evasion of the immune response^7^; the mutation N501Y, which enhances binding affinity for the human ACE2 receptor and therefore influences viral transmissibility^8, 9^; and the mutation P681H, which is adjacent to the S1/S2 furin cleavage site in spike and might have an impact on viral infectivity^10, 11^.

A recent study indicated that the SARS-CoV-2 VOC carrying the 501Y mutation showed no higher infectivity in cell than ancestral D614G variants^12^. Likewise, we did not observe replication of the B.1.1.7 variant to be significantly enhanced over that of other tested variants at any of the selected time-points in Vero-E6 and Calu-3 cells (Extended Data Fig. 1), however, we did observe that B.1.1.7 dominates in competitive fitness assays. These comparisons of replication fitness between B.1.1.7 and earlier circulating strains were performed in Calu-3 cells through simultaneous co-infection at a 1:1 ratio with B.1.1.7 (accession number: MW856794) and another variant of the D614G lineage, either B.1-G (HK-95, accession number: MT835143) or B.1.GH (405, accession number: MW856793) (Fig. 1). After three rounds of consecutive passage at 72-hour intervals, the B.1.1.7 variant became dominant in both co-culture conditions, suggesting that the additional substitutions in B.1.1.7 enhance SARS-CoV-2 replication fitness in cells.

**Figure 1.**
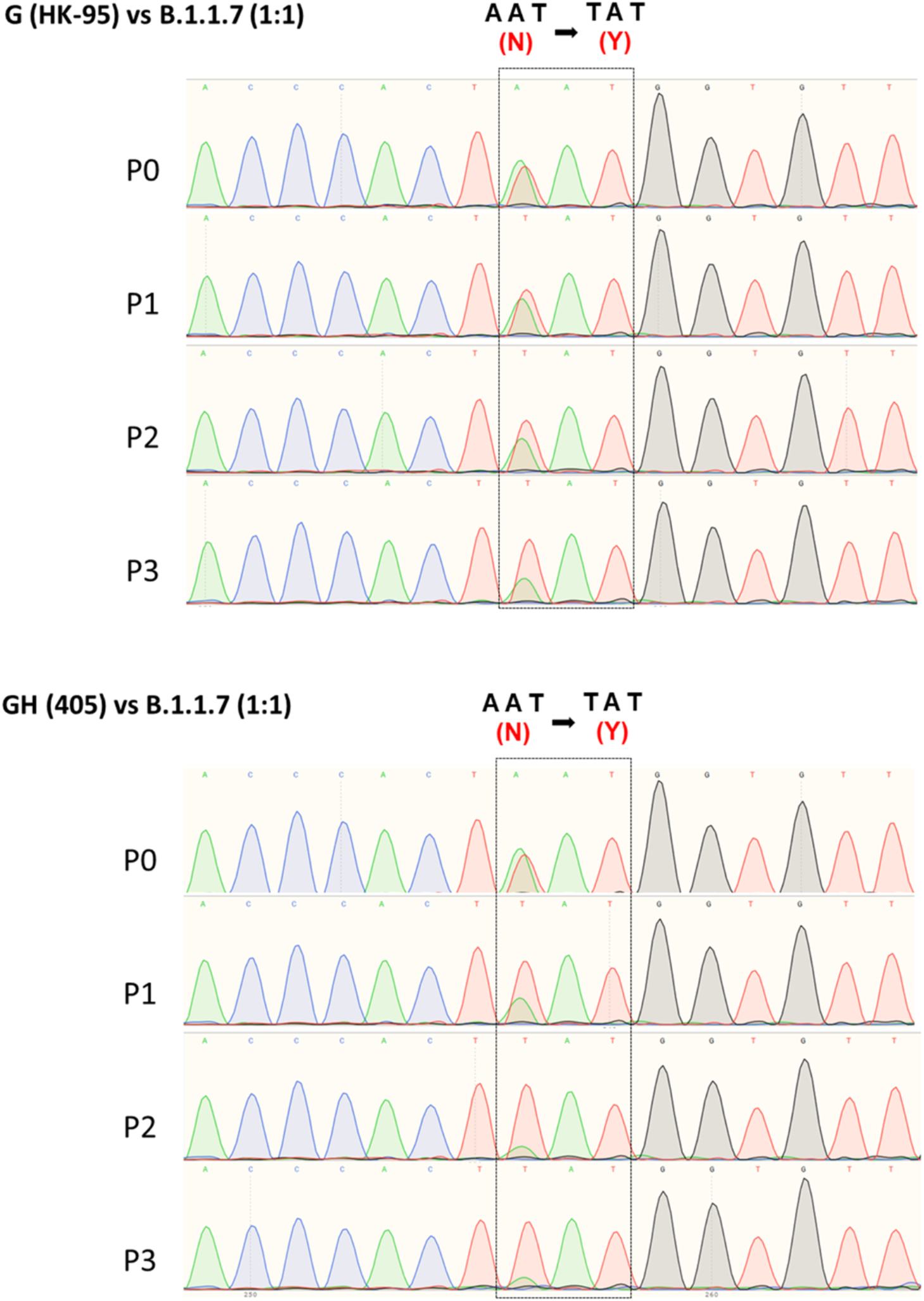
*In-vitro* Competitive Fitness Assay. Sanger sequencing chromatograms of spike gene fragments amplified from viral samples in the competition assay. Cell cultures were infected with a 1:1 mixture of two variants, as indicated, at an MOI of 0.1. The supernatants were serially passaged three times in Calu-3 cells. 901 bp fragments containing residue 501 (boxed) were amplified from the vRNA of individual samples collected from each passage (P) and sequenced. B.1-G (HK-95) and B.1.GH (405) are 501N, B.1.1.7 is 501Y.

Next, we set up a Syrian hamster infection study to evaluate if B.1.1.7 exhibits higher infectivity *in vivo*. 6-8-week-old male Syrian hamsters were intranasally infected with 50 microliters of different variants (2×10^4^ PFU/ml), which is equivalent to 1000 PFU per inoculum, as indicated in Fig. 2A. Infectious viral titres in upper (nasal) and lower (pulmonary) tissues were measured on four consecutive days after infection. All viruses tested replicated to similar titres in nasal turbinate and lung tissues of infected hamsters. This result is consistent with two recent studies which also found no significant alteration in infectious viral titres in samples collected from nasal washes, throat swabs and lungs from hamsters infected with different SARS-CoV-2 variants^13,14^. Given that hamsters are highly susceptible to SARS-CoV-2 infection, intranasal infection with high-titre inocula may hamper discrimination of differences in the infectivity and replication efficiency of variants^15^. In fact, by titrating the infection dosage (10-fold dilution) of the inocula administered to hamsters, we observed that viral replication in nasal tissues of infected hamsters had already plateaued with infection doses of 100 PFU and upwards, even on day one post-infection (Extended data Fig. 2). Humans are exposed to varying doses of infectious particles during SARS-CoV-2 transmission. We reasoned that SARS-CoV-2 variants which can initiate effective infection with fewer infectious particles are likely to transmit more effectively than other variants requiring more infectious particles. To test this, we performed another hamster infection study using only 10 PFU per inoculum, with samples being collected at 16 hours post-infection. Interestingly, infectious viral loads in nasal turbinates of hamsters were found to be significantly higher with B.1.1.7 compared to the other viruses, whereas similar viral loads were observed in lungs of all infected hamsters, except for those inoculated with B.1-G (HK-95), which exhibits higher viral titres in lungs, although with large variations between replicates (Fig. 2B).

**Figure 2.**
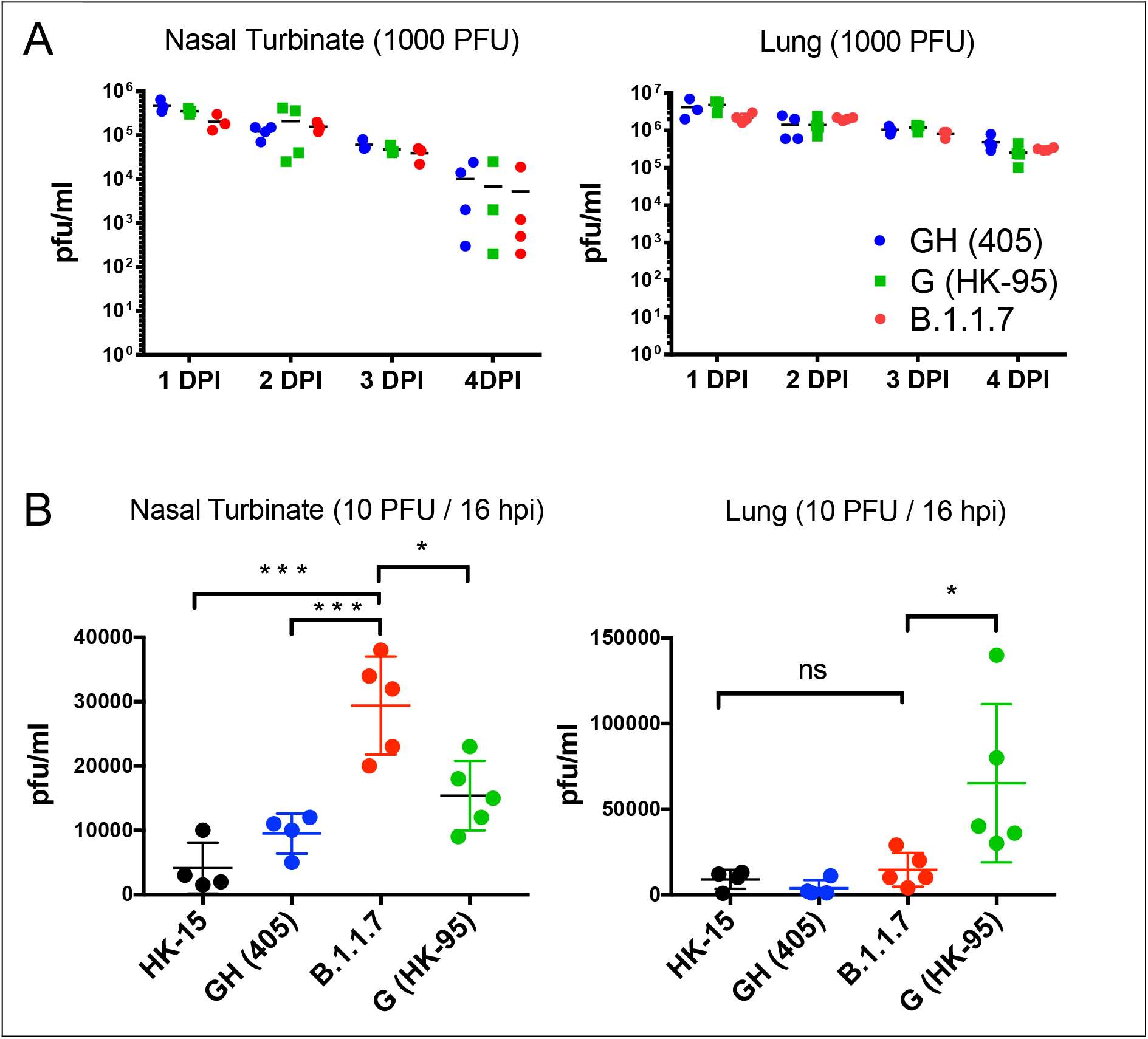
*In-vivo* Infection Studies. Viral replication of different SARS-CoV-2 variants in nasal turbinates and lungs of hamsters. Hamsters were infected with different SARS-CoV-2 variants, as indicated. Viral titers in nasal turbinates and lungs were determined by plaque assay (PFU/ml). (A) Hamsters (14 per variant virus group) were each inoculated intranasally with 50 ul of virus stock containing 1000 PFU of virus. Three to four hamsters from each group were euthanized on each of the four consecutive days following infection for viral titration. (B) Hamsters (4-5 per group) were each inoculated intranasally with 50 ul of virus stock containing 10 PFU of virus. One non-D614G lineage variant (HK-15 (MT835141)) and three D614G lineage variants (GH (405), B.1.1.7 and G (HK-95)) were used. Hamsters were euthanized at 16 hours post-infection for viral titration. Horizontal lines indicate the overall mean of average viral titer values per group. Statistical significance was calculated by Student’s t-test; * denotes *p<0*.*05*, *** denotes *p<0*.*0005* and ns denotes *non-significant*.

SARS-CoV-2 VOCs have been emerging in different countries in the past few months, and it is crucial to establish relevant experimental models to characterise existing and new variants in terms of transmissibility, disease severity and vaccine efficacy, and to evaluate therapeutic interventions. In this report, by using a lower infectious dose, we demonstrate that B.1.1.7 exhibits higher infectivity and/or replication efficiency in the nasal epithelium. Our data, albeit limited, strengthen the contention that this novel VOC is more easily transmitted than other pre-existing strains. Further work, including transmission studies with optimised inoculum dosages and timing of sample collection and investigation into routes of transmission are required. A better understanding of SARS-CoV-2 dynamics is important for designing combative strategies for the prevention and control of virus infections.

## Acknowledgments

The authors would like to thank Dr Jane Rayner for critical reading and editing of the manuscript. This study is partly supported by the Theme-Based Research Scheme (T11/707/15) and General Research Fund (17107019) of the Research Grants Council, Hong Kong Special Administrative Region, China, and the Sanming Project of Medicine in Shenzhen, China (No. 290 SZSM201911014).

## Declaration of interest statement

No potential conflict of interest was reported by the author(s).

## Methods

### Viruses

The SARS-CoV-2 isolates HK-95 (MT835143), 405 (MW856793), B.1.1.7 (MW856794) and HK-15 (MT835141) were isolated from specimens obtained from four laboratory-confirmed COVID-19 patients using Vero E6 cells (ATCC; CRL-15786). All experiments involving SARS-CoV-2 viruses were conducted in a Biosafety Level-3 laboratory. For animal challenge, viral stocks were prepared after two serial passages of isolated virus in Vero E6 cells in Dulbecco’s Modified Eagle Medium (DMEM) (Thermo Fisher Scientific) supplemented with 5% fetal bovine serum (Thermo Fisher Scientific), and 100 IU penicillin G/ml and 100 ml streptomycin sulfate/ml (Thermo Fisher Scientific). Virus titres were then determined by plaque assay using Vero E6 cells. Viral RNAs were obtained from the supernatants of infected cells and then isolated using the QIAamp RNA Viral kit (Qiagen) and subjected to whole viral genome sequencing.

### Hamster Infection

Female golden Syrian hamsters, aged 6-8 weeks old, were obtained from the LASEC, Chinese University of Hong Kong via the Centre for Comparative Medicine Research at the University of Hong Kong (HKU). All experiments were performed in a Biosafety Level-3 animal facility at the LKS Faculty of Medicine, HKU. All animal studies were approved by the Committee on the Use of Live Animals in Teaching and Research, HKU. Hamsters were anesthetized with ketamine (150mg/kg) and xylazine (10 mg/mg) via intraperitoneal injection prior to nasal inoculation. All hamsters were euthanized by intraperitoneal injection of pentobarbital at 200 mg/kg.

### *In-vitro* Competitive Fitness Assay

Calu-3 cells in Dulbecco’s Modified Eagle Medium (DMEM) (Thermo Fisher Scientific) supplemented with 5% fetal bovine serum (Thermo Fisher Scientific), and 100 IU penicillin G/ml and 100 ml streptomycin sulfate/ml (Thermo Fisher Scientific) were infected with MOI of 0.1 of B.1.1.7 and another variant of the D614G lineage, either B.1-G (HK-95) or B.1.GH (405) mixture at 1:1 ratios. Following 1h incubation, the cultures were washed thrice with PBS and cultures for 3 days. To passage the progeny viruses, the virus samples were continuously passaged three times in Calu-3 cells. Viral RNAs were obtained from the supernatants of infected cells and then isolated using the QIAamp RNA Viral kit (Qiagen). A 901 bp fragment containing the N501Y site was amplified from each RNA sample by RT-PCR using primer set: 5’-GAAGTCAGACAAATCGCTCCAG-3’ and 5’-GCAACTGAATTTTCTGCACCA-3’. The amplicon was purified by NucleoSpin® Gel and PCR Clean-Up (Takara) for Sanger sequencing.

## Notes

### Competing Interest Statement

The authors have declared no competing interest.

