## Extended data for "Low dose inocula of SARS-CoV-2 B.1.1.7 variant initiate more robust infections in the upper respiratory tract of hamsters than earlier D614G variants"

**Extended Data Fig.1.** **Growth curves of B.1.1.7, 405 and HK-95 in Calu-3 cells at a MOI = 0.01.** Virus infected cells were cultured at 37°C and supernatants then harvested at the indicated time points and subjected to plaque assay in Vero E6 cells to determine the virus titre.

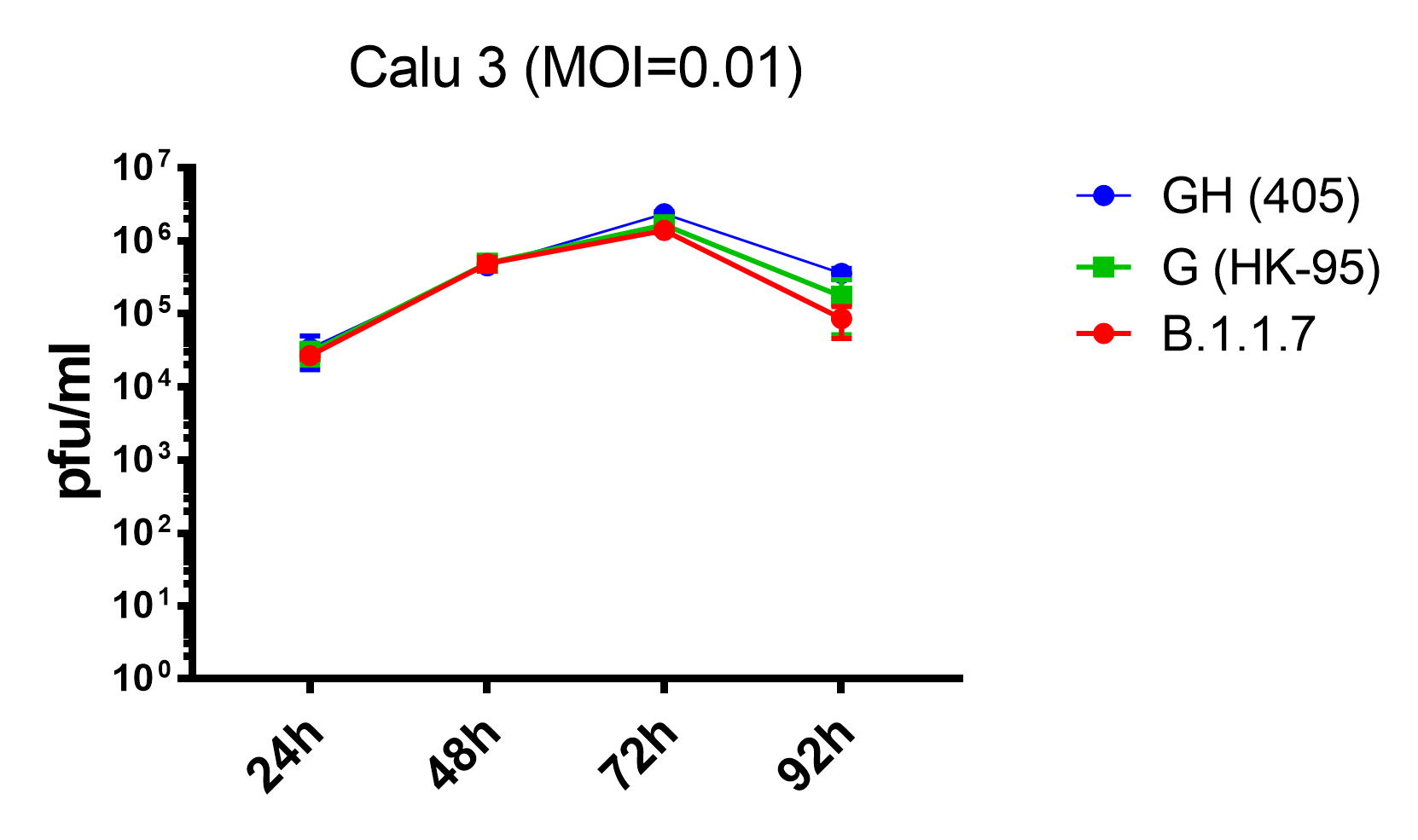

### Extended Data Fig.2. Viral replication in nasal turbinates and lungs of hamsters infected with different dose inocula of B.1.1.7. Hamsters (2 per group) were each inoculated intranasally with different dose inoculums of B.1.1.7 as indicated. All hamsters were euthanized on one day post-infection for viral titration. Horizontal lines indicate the overall mean of average viral titer values per group. Statistical significance was calculated by Student’s t-test; * denotes *p<0.05*.

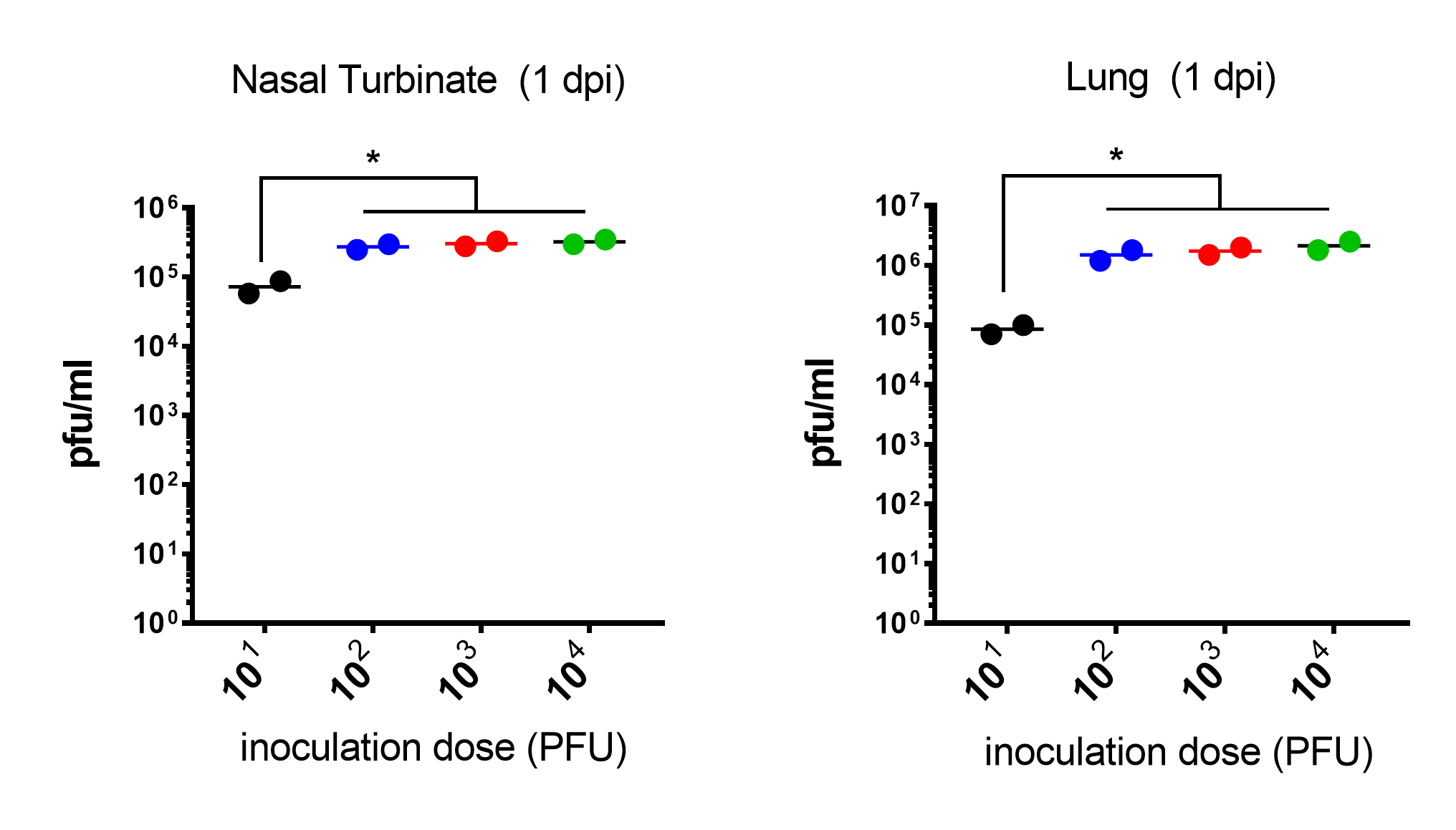
